# Probability Ramp Control reduces the number of sessions required to find an acceptable dose of succinylcholine during Electroconvulsive Therapy - an *in silico* analysis

**DOI:** 10.1101/2022.08.04.502847

**Authors:** Jeff E Mandel

## Abstract

**Introduction:** Electroconvulsive Therapy may be utilized in as many as 76,000 cases annually in the US, with the majority of cases employing succinylcholine. The reported dose spans the range of 0.29 - 2.1 mg/kg, and while motor seizures only last 36 ± 6 seconds, the duration of paralysis extends to 310 ± 38 seconds. While a model of succinylcholine pharmacokinetics/pharmacodynamics exists, this has not been employed to facilitate dose selection in clinical practice. Probability ramp control was investigated for this purpose.

**Methods:** Two approaches to dose finding were implemented. The first was an optimized Up-Down Method (UDM) that utilized an initial bolus, an adjustment dose, and a decrement to decrease the adjustment when crossing the target of 95% twitch depression. The second utilized probability ramp control (PRC) comprised of an infusion sequence that stopped when 95% twitch depression was obtained, a model that mapped the times for onset and offset of blockade to a subsequent bolus, and an adjustment dose to refine this dose when needed. Two populations of 10000 randomly parameterized models were developed from published data to train and evaluate the performance. Performance was assessed with a fuzzy classifier that segmented outcomes into three sets – LOW, HIGH, and SUCCESS. A loss function was developed that determined the number of sessions required to bring all models to SUCCESS. The probability distributions were compared using the Kolmogorov-Smirnov 2 sample test, with P<0.001 considered significant.

**Results:** Optimal values for the UDM parameters BOLUS, ADJUSTMENT, and DECREMENT were 0.7834 mg/kg, 0.3334 mg/kg, and 0.4056. Optimal values for the PRC SEQUENCE were 0.2663 mg/kg/min for 3 minutes followed by 0.7028 mg/kg/min. A fourth order polynomial MODEL produced estimates of the bolus that brought 99% of models to SUCCESS on the second session, while UDM required 6 sessions to achieve 99% SUCCESS. The probability distributions were distinct with P<<0.001.

**Discussion:** PRC was able to correctly produce SUCCESS in significantly fewer sessions than UDM. Additionally, PRC is easy to implement and allows pooling of results from multiple clinicians. The performance of PRC in clinical use for ECT will require further study.

**Key Points:** 

**Question:** Can probability ramp control reduce the number of ECT sessions with suboptimal succinylcholine dosing?

**Findings:** Probability ramp control found the correct dose in two sessions in 99% of simulations, compared to six sessions for the Up-Down Method.

**Meaning:** Probability ramp control is a more efficient method for finding the appropriate dose of succinylcholine for repeated sessions of ECT.

## Introduction

Electroconvulsive therapy (ECT) is widely practiced technique for treatment of refractory depression. Most ECT utilizes neuromuscular blocking agents, typically succinylcholine. Succinylcholine has considerable variability in pharmacokinetics and pharmacodynamics. In a retrospective review of 500 patients Bryson^1^ found succinylcholine doses span the range of 0.29 – 2.1 mg/kg, with 180 patients requiring an adjustment from the standard dose of 0.9 mg/kg. The duration of paralysis from succinylcholine may exceed the requirement for prevention of movement during the motor seizure. In a prospective comparison of succinylcholine to rocuronium for ECT, Kadoi^2^ found a dose of 1 mg/kg produced an average duration of > 90% twitch reduction of 310 ± 38 seconds, while motor seizure duration was only 36 ± 6 seconds. While Dixon’ s Up-and-Down method (UDM) has been applied to muscle relaxant dose finding in ECT^3^, this only established that 0.85 mg/kg of succinylcholine produces > 95% twitch reduction in 50% patients. Since there are adverse consequences to both inadequate control of motor seizures and to prolonged paralysis, finding the correct dose for the individual patient in the minimum number of sessions would seem useful.

Recall of paralysis is unpleasant, and may cause some patients to refuse further ECT treatments. To avoid recall of paralysis, sedative agents such as methohexital are employed. Because these agents interfere with seizure production, there is a limit on the size of bolus of sedative that can be utilized and the duration of amnesia this dose will produce. A dose of succinylcholine that yields paralysis exceeding the duration of motor seizure but less than the duration of amnesia is acceptable; finding a search strategy that minimizes the number of ECT sessions with unacceptable results is the goal. This notion can be expressed as a loss function that considers the number of sessions which fail to achieve 95% twitch reduction for one minute and recovery within 6 minutes.

Probability ramp control (PRC) is an approach to dose finding that uses prior information to increase the drug concentration monotonically through the pharmacodynamic area of interest over a specified time course.^4^ The method requires an initial estimate of the probability density function for the intended effect of succinylcholine. Roy^5^ developed a combined pharmacokinetic-pharmacodynamic model of succinylcholine, which was utilized to develop this probability density function from a bootstrapped population. I hypothesized that PRC would produce fewer suboptimal ECT sessions than optimized UDM.

## Methods

The Roy model was implemented in state-space form in MATLAB 2021b (Mathworks, Natick, MA) and each of the model parameters was fit to a gamma distribution to permit generation of random models for bootstrap analysis. Note that the gamma distribution was utilized to avoid generating models with non-positive model parameters. Models were assessed to determine the dose required to achieve a one minute duration of 95% twitch depression and the dose required to achieve recovery to 95% twitch reduction at six minutes; models yielding a dose outside the range of 0.29 - 2.1 mg/kg were excluded from further analysis. Source code is included in the supplemental material.

A loss function was implemented by simulating the trajectory for the succinylcholine effect site concentration for the prescribed administration and computing both the time above 95% twitch reduction and the time at which recovery below 95% twitch recovery occurred. This trajectory was used to form a fuzzy set; if the duration of paralysis was less than one minute, the score was LOW; if recovery was not seen in six minutes, the score was HIGH; otherwise, the score was SUCCESS. Examples of the fuzzy score are depicted in Figure 1. The loss function was obtained by determining the number of sessions required to bring all models to SUCCESS.

**Figure 1:**
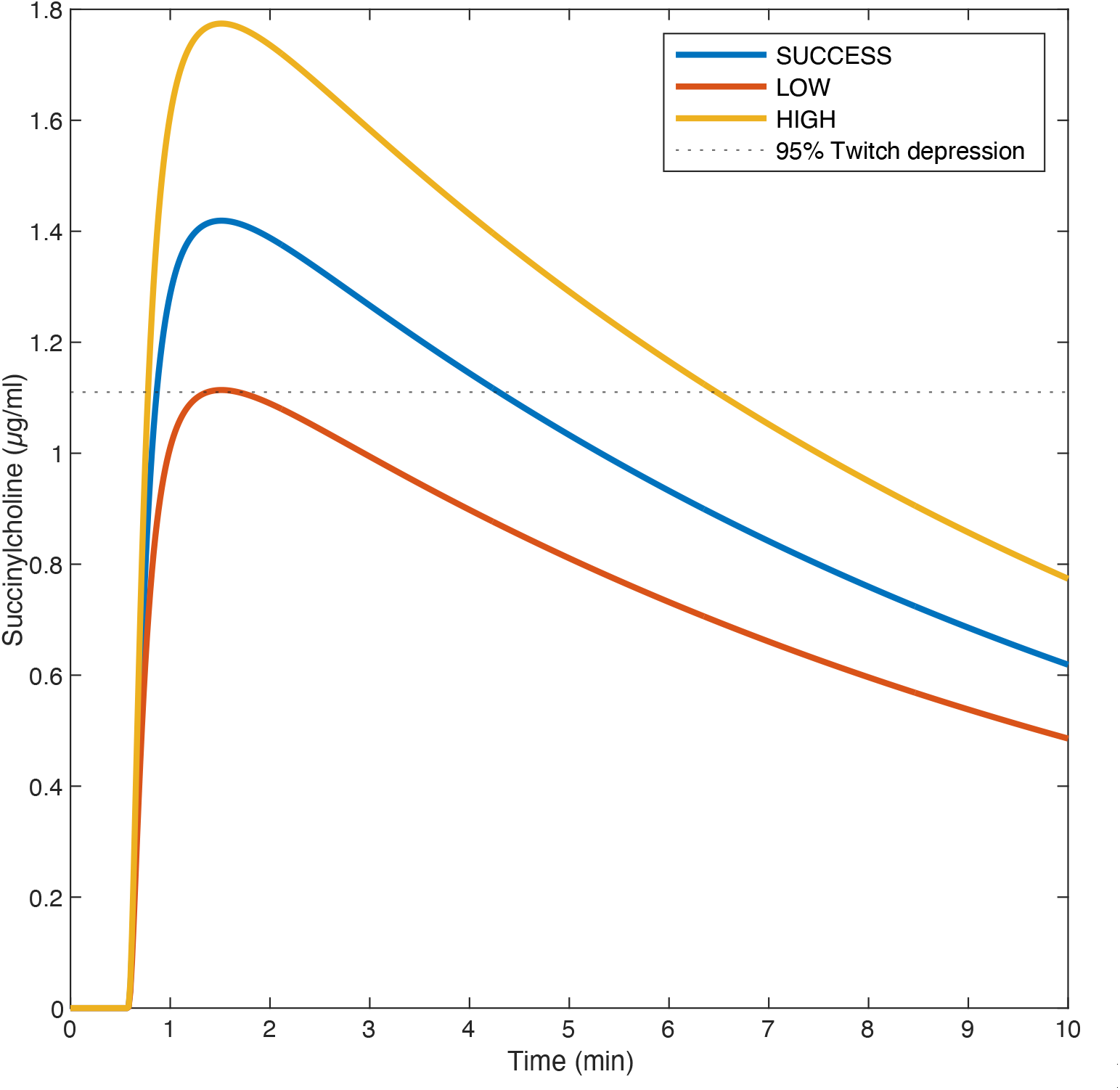
Trajectory of three boluses applied to a single model, illustration LOW (red), HIGH (yellow), and SUCCESS (blue).

Two populations of simulated patients were generated using distinct random seeds. The first population of 10,000 patients was used for determining optimal parameters of the UDM and PRC, the second for determining the value of the loss function with these settings. These are referred to as “training” and “scoring”.

The UDM was configured with three parameters - BOLUS, ADJUSTMENT, and DECREMENT. The BOLUS was administered to each model in the first session and the resulting fuzzy score determined. If the score on the previous session changed from LOW to HIGH or HIGH to LOW, ADJUSTMENT was decreased by DECREMENT. If score was LOW, bolus was increased by ADJUSTMENT, if HIGH, BOLUS was decreased by ADJUSTMENT. Once the BOLUS yielded SUCCESS, bolus adjustment ceased. This is depicted in Figure 2. The values for BOLUS, ADJUSTMENT, and DECREMENT were obtained using simplex minimization to minimize the loss function across the training model set. These optimized parameters were then applied to the scoring model set to determine the number of sessions required to bring all models to SUCCESS.

**Figure 2:**
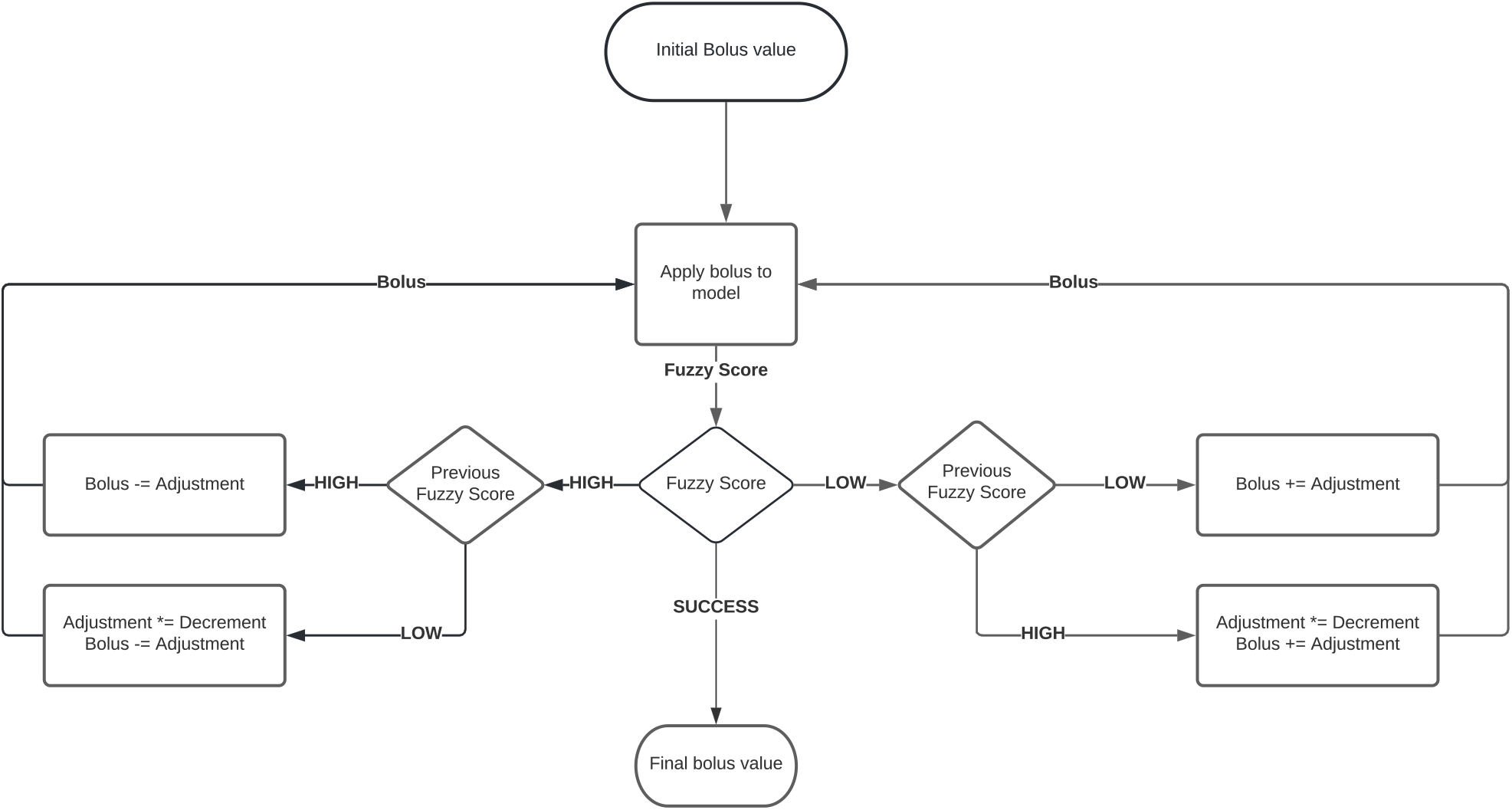
Flowchart illustrating the Up-Down Method

The PRC was configured with three parameter sets – SEQUENCE, MODEL, and ADJUSTMENT. SEQUENCE is an infusion sequence comprised of an initial rate delivered over three minutes followed by a secondary rate delivered over the subsequent three minutes. This sequence was applied to each training model and the duration required to achieve one minute above 95% twitch depression determined. An example of this process is depicted in Figure 3. A loss function that compared this collection of durations (ESTIMATED) to a normal distributed of time with a mean of three minutes and a standard deviation of one minute (TARGET) was used to obtain the optimal settings for SEQUENCE, as shown in Figure 4. This infusion sequence was applied to each training model for the first simulated session until 95% twitch depression was observed, when the infusion sequence was terminated, and the time to recovery to 95% twitch depression was determined. MODEL is a model estimated by linear regression to predict the optimal bolus from these aforementioned times, and is used to determine the starting bolus for the second session. If a third session is required, ADJUSTMENT is used to modify the bolus; separate values for the LOW and HIGH cases were determined. These optimized parameter sets were applied to the scoring model set to determine the number of sessions required to bring all models to SUCCESS.

**Figure 3:**
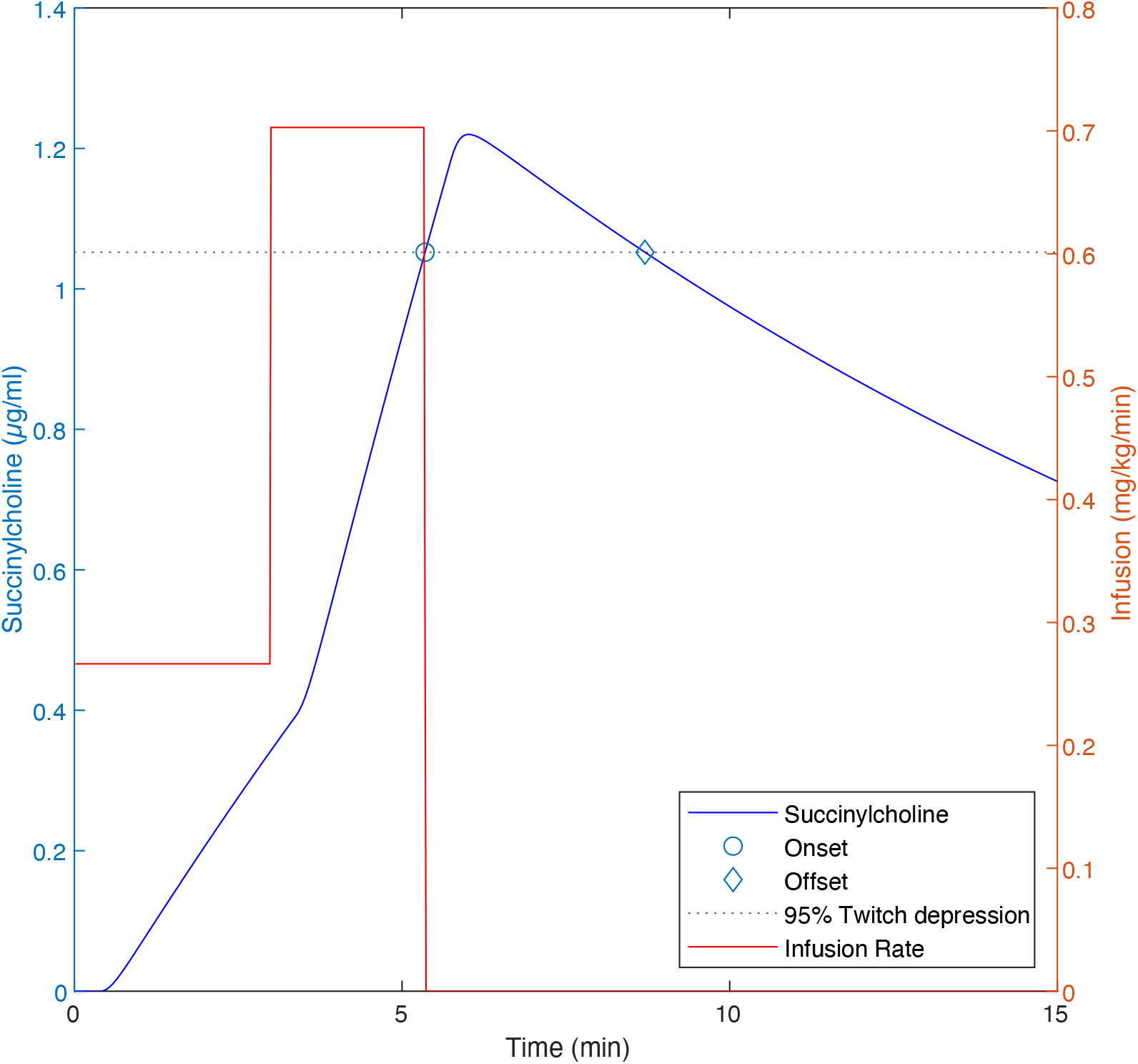
Application of PRC to a single model. The infusion rate of 0.2663 mg/kg/min proceeds for 3 minutes, then increases to 0.7028 mg/kg/min for 2.36 minutes, when 95% twitch depression is observed and the infusion is stopped, having delivered a total of 2.46 mg/kg. Recovery to 95% twitch depression occurs at 8.71 minutes. The optimal bolus for this model is 2.27 mg/kg.

**Figure 4:**
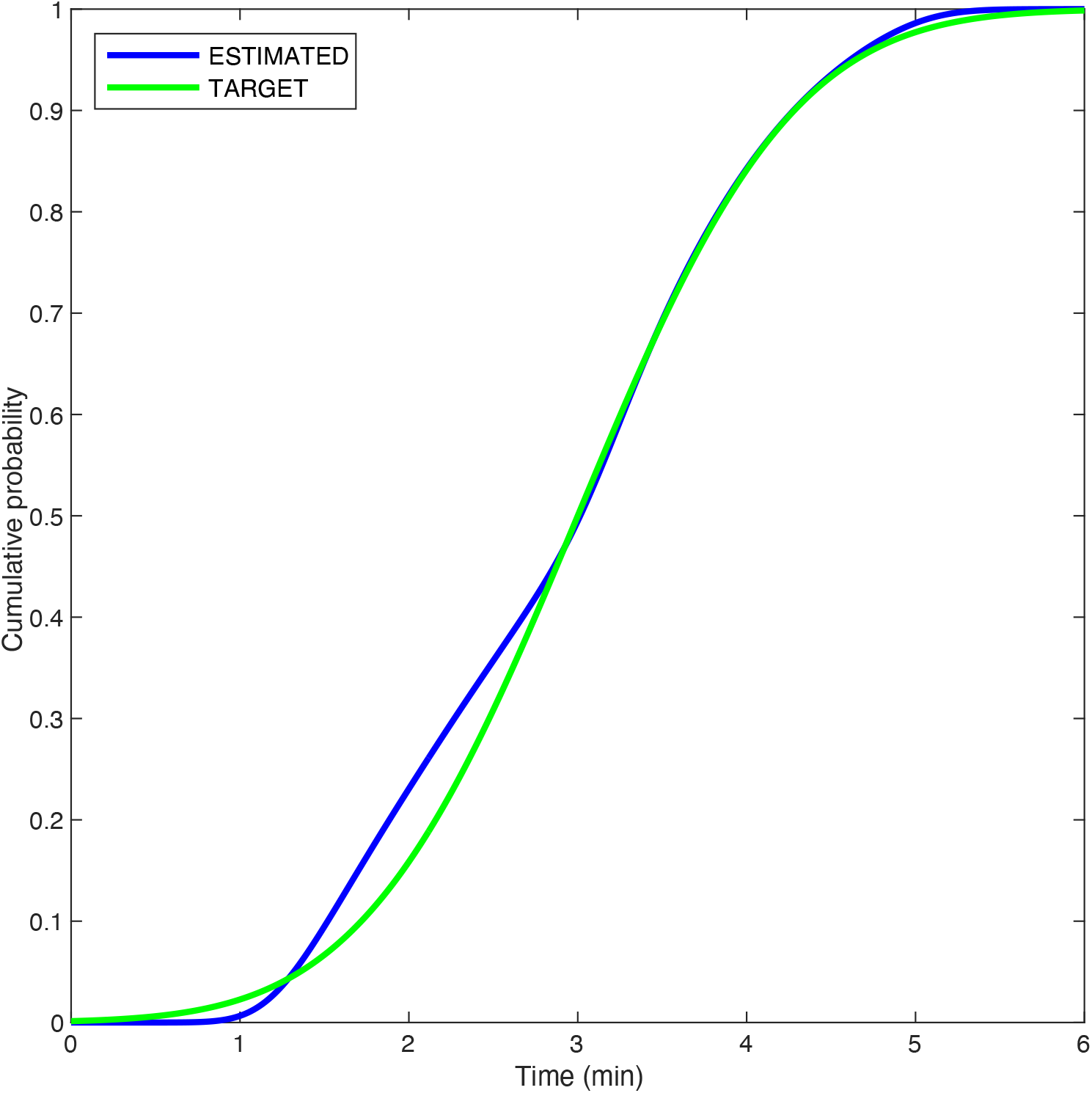
The temporal objective function for Probability Ramp Control, illustrating TARGET (a normally-distributed time of mean 3 minutes and standard deviation 1 minute, green) and the result of applying the optimal sequence to the training model set (blue).

The number of sessions required to bring all models to SUCCESS for the two groups were compared by the Kolmogorov Smirnov 2 sample test. P<0.001 was considered significant.

## Results

The median dose required to achieve one minute of 95% twitch depression was 0.77 mg/kg. The median dose yielding 95% twitch recovery in six minutes was 0.94 mg/kg.

Optimal values for the UDM parameters BOLUS, ADJUSTMENT, and DECREMENT were 0.7834 mg/kg, 0.3334 mg/kg, and 0.4056. Optimal values for the PRC SEQUENCE were 0.2663 mg/kg/min and 0.7028 mg/kg/min. The PRC parameter MODEL was estimated using a polynomial model with increasing terms. A fourth order polynomial yielded an R^2^ of 0.994 with a MSE of 0.039, which was sufficient to obtain SUCCESS in 99% of models on the second session. ADJUSTMENT was 0.0549 mg/kg for LOW and 0.0704 mg/kg for HIGH.

A histogram of the number of sessions required to bring all models to SUCCESS in the two groups is presented in Figure 5. The distributions of the two groups were different with P << 0.001. Of note, while on the initial session 14% of the UDM models vs. 3.3% of the PRC models met the criteria for SUCCESS, on the second session PRC attained SUCCESS in 99% of models; UDM did not achieve this level of SUCCESS until the 6^th^ session. In the first session of PRC all models not in SUCCESS were in HIGH, while in UDM, 95% of models not in SUCCESS were in LOW.

**Figure 5:**
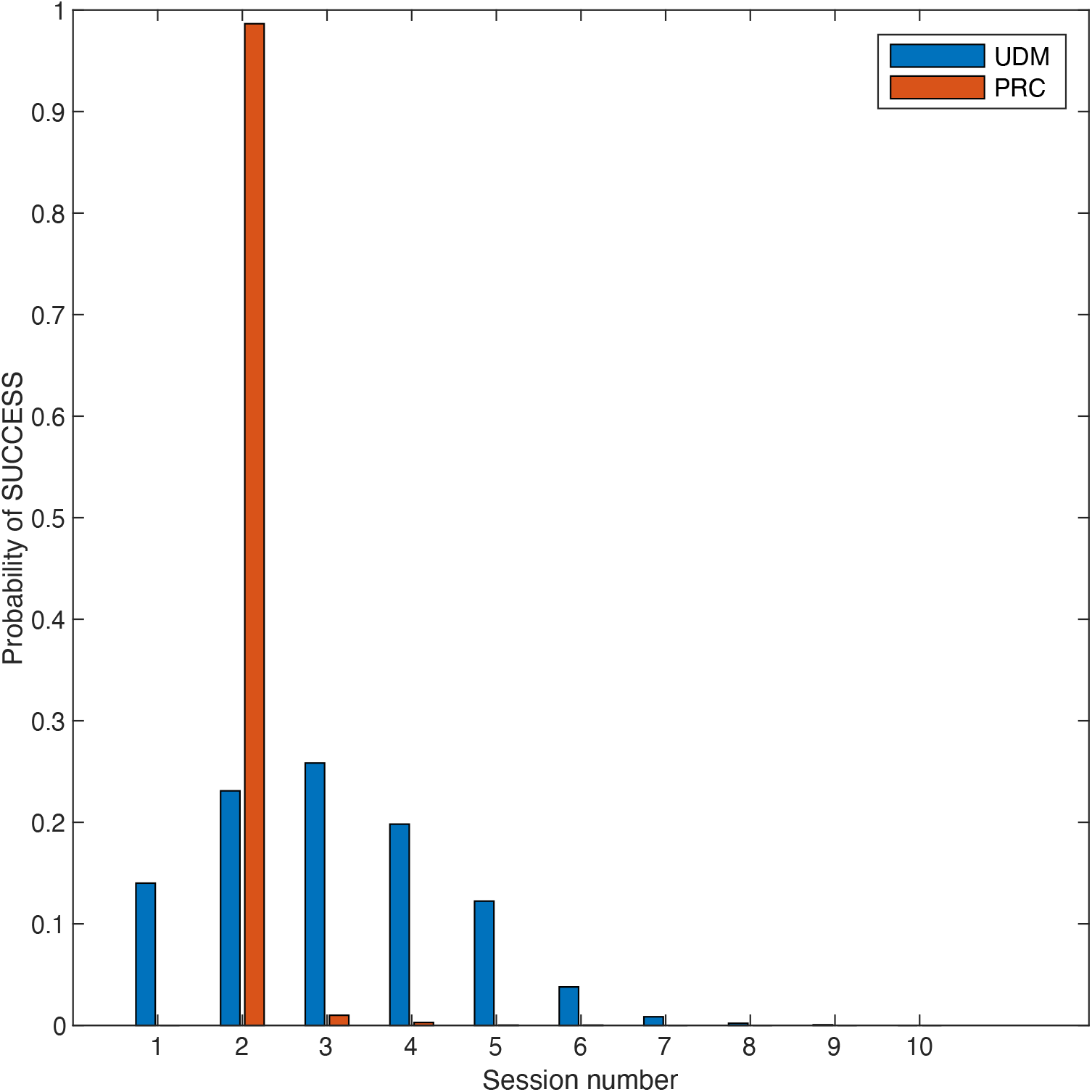
Histogram of the number of models meeting SUCCESS in UDM (blue) vs. PRC (red)

## Discussion

The most important finding of this effort is that use of PRC to drive administration of succinylcholine in the initial session of ECT can permit precise prediction of bolus dosing for subsequent sessions. While the optimized UDM was able to achieve SUCCESS in a larger number of initial sessions than PRC, a 14% rate of SUCCESS is insufficient to obviate a backup strategy for incomplete paralysis. While such a strategy could consist of giving an additional dose of succinylcholine when 95% paralysis has not been observed after two minutes, this would frequently extend the recovery beyond six minutes. In the model depicted in Figure 3, seven additional ADJUSTMENT boluses given at two minute intervals (for a total of 3.12 mg/kg) would be required to achieve 95% twitch reduction. In PRC a strategy to prevent awareness of residual paralysis on the initial session is required. Such a strategy could be a larger induction dose of sedative, subsequent boluses of sedative, or a continuous infusion of sedative after the initial bolus. An advantage to employing PRC in the dose finding process is that the information needed to predict the dose for the second session is available at the conclusion of the first session, while in UDM this information may not be available until the bolus produces SUCCESS. This may make the process of initiating therapy in a new patient more efficient.

PRC is designed to be easily implemented with commonly available volumetric pumps. An initial bolus of 0.8 mg/kg delivered over three minutes followed by an infusion of 0.7028 mg/kg/min is all that is needed to perform the initial titration. While an automated system for assessing neuromuscular blockade may be useful, a simple peripheral nerve stimulator is likely to be adequate. The calculation of bolus dosing for the second and subsequent sessions can be performed with a smart phone or web app; a lookup table for deciles of onset and offset times is presented in Table 1.

**Table 1.** Lookup table to determine estimated bolus (mg/kg) from the onset and offset times (min) for the PRC load.

|  |  | Onset time (min) |  |  |  |  |  |  |  |  |
| --- | --- | --- | --- | --- | --- | --- | --- | --- | --- | --- |
|  |  | 2.1 | 2.42 | 2.75 | 3.11 | 3.47 | 3.75 | 4.02 | 4.33 | 4.75 |
| Offset time (min) | 7.02 | 0.48 | 0.56 | 0.67 | 0.83 | 1.03 | 1.22 | 1.43 | 1.68 | 2.04 |
|  | 8.07 | 0.46 | 0.53 | 0.63 | 0.78 | 0.97 | 1.15 | 1.34 | 1.59 | 1.92 |
|  | 8.89 | 0.44 | 0.51 | 0.61 | 0.75 | 0.93 | 1.1 | 1.29 | 1.53 | 1.85 |
|  | 9.69 | 0.43 | 0.5 | 0.59 | 0.73 | 0.9 | 1.07 | 1.25 | 1.48 | 1.79 |
|  | 10.43 | 0.42 | 0.49 | 0.58 | 0.71 | 0.88 | 1.04 | 1.22 | 1.44 | 1.75 |
|  | 11.26 | 0.41 | 0.48 | 0.56 | 0.69 | 0.86 | 1.01 | 1.19 | 1.4 | 1.71 |
|  | 12.29 | 0.39 | 0.46 | 0.55 | 0.67 | 0.83 | 0.99 | 1.15 | 1.36 | 1.66 |
|  | 13.59 | 0.38 | 0.45 | 0.53 | 0.65 | 0.81 | 0.96 | 1.12 | 1.33 | 1.62 |
|  | 15.53 | 0.36 | 0.43 | 0.51 | 0.63 | 0.78 | 0.92 | 1.08 | 1.28 | 1.57 |

Luccarelli ^6^ utilized freedom of information filings to estimate utilization of ECT in US states with mandated reporting, and found a range of 0.5 – 2.4 patients/10,000 residents, suggesting between 16,000 and 76,000 cases annually in the US. Use of PRC might eliminate suboptimal succinylcholine dosing in a significant fraction of these patients. Additionally, wide utilization of PRC with central registry could rapidly provide guidance on dosing of this drug based on objective clinical observations. While this effort is concerned with management of patients undergoing ECT, there are other paradigms in which anesthesiologists care for patients on multiple occasions, particularly in non OR anesthesia settings. The availability of tools to provide consistency in the management of such procedures may be useful.

This study has an important limitation – the sample population was derived from seven patients. Despite this, the median doses for one minute of paralysis and recovery at six minutes are in fair agreement with the observations of clinical studies, and are unlikely to favor one method or the other. Probability Ramp Control is not dependent on the accuracy of the effect site concentration predictions; it deals with the mapping of drug delivery to observed clinical effects. While an initial estimate of the probability distribution for this mapping is required, once a sufficient number of patients have been observed, the actual probability distribution can replace the hypothetical distribution. This mimics the behavior of clinicians – we begin with only “book knowledge”, but slowly develop clinical experience. Clinical experience is not easily shared, and systems that rely on a small number of clinicians with expertise in a niche practices may not scale well. Probability ramp control offers the potential to merge observations of many clinicians to answer questions not easily addressed by conventional pharmacokinetic/pharmacodynamic methods. The utility of PRC in optimizing dose selection of succinylcholine during ECT will require clinical validation.

## Supporting information

Supplemental Code listing

## Glossary of terms

ECT: Electroconvulsive therapy
PRC: Probability ramp control
UDM: Up-Down Method

## Notes

### Competing Interest Statement

The authors have declared no competing interest.

