## Supplemental Code listing for "Probability Ramp Control reduces the number of sessions required to find an acceptable dose of succinylcholine during Electroconvulsive Therapy - an *in silico* analysis"

```
function [models,oneminute_target, max_target] = succinylcholine(numModels, seed)
% Implements the model described in:
% Roy JJ, Donati F, Boismenu D, Varin F. Concentration-effect relation of
% succinylcholine chloride during propofol anesthesia. Anesthesiology. 2002
% Nov;97(5):1082-92. doi: 10.1097/00000542-200211000-00009. PMID: 12411790.
%
% Returns a cell array of numModels randomly distributed models. If seed is
% passed, use this to initialize the random number generator, otherwise use
% default.
%
% 2021 Jeff E Mandel MD MS

if nargin == 2
    rng(seed);
else
    rng('default');
end
% weights (kg)
weights = [80;71;56;80;88;77;68];

% Microrate constants (min^-1)
k01 = [10.71;5.55;11.64;14.29;6.54;25.00;30.84];
k10 = [2.2;2.4;6.3;5.7;3.3;5.4;9.7];
k12 = [0.5;1.1;2.3;1.3;0.9;2.6;1.7];
k21 = [1.0;1.1;1.7;1.9;1.6;1.7;1.9];
k20 = [0.74;0.67;1.21;1.45;1.12;1.05;0.82];

% delay (minutes)
delay = [0.53;0.15;0.38;0.44;0.43;0.36;0.29];

% Volumes (ml/kg)
V1 = [11;12;6;9;13;7;4];
% Vss = [17;24;14;15;20;17;28];
% Vsscp = [34;45;34;36;41;35;48];

%PD parameters
ke0 = [0.0318;0.03804;0.06948;0.0327;0.06774;0.07476;0.09474];
gamma = [18.6;27.1;12.4;33.7;15.5;14.0;13.5];
EC50 = [.630;.671;.855;.397;.788;.995;.970];

%Parameterize twice as many as we need so that we can discard ones that
%fall outside the specified range
tempModels=round(numModels*2);
weight = mean(weights);
k01r = get_random(k01, tempModels);
k10r = get_random(k10, tempModels);
k12r = get_random(k12, tempModels);
k21r = get_random(k21, tempModels);
k20r = get_random(k20, tempModels);
V1r = get_random(V1, tempModels).*weight;
delayr = get_random(delay, tempModels);

ke0r = get_random(ke0, tempModels);
gammar = get_random(gamma, tempModels);
```

```
EC50r = get_random(EC50, tempModels);

models = cell(numModels,1);
max_target=nan(numModels,1);
oneminute_target=nan(numModels,1);
target = 0.95;

B = [weight;0;0;0];
D = 0;
h1 = waitbar(0);
i=1;
k=1;
while i<=numModels
    waitbar(i/numModels,h1);
    k01a = k01r(k);
    k10a = k10r(k);
    k12a = k12r(k);
    k21a = k21r(k);
    k20a = k20r(k);
    ke0a = ke0r(k);
    V1a = 1E3./V1r(k);
    A = [-k01a 0            0          0
          k01a -(k10a+k12a) k21a       0
          0       k12a      -k20a-k21a 0
          0       ke0a      0          -ke0a];
    C = [0 0 0 V1a];
    model.sys = ss(A,B,C,D,'InputDelay', delayr(k));
    model.gamma = gammar(k);
    model.EC50 = EC50r(k);
    model.target_conc = findConc(model, target);
    [oneminute,max] = modelTargets(model);
    if oneminute<=2.1 && oneminute>= 0.29 && oneminute<max
        model.oneminute_target = oneminute;
        model.max_target = max;
        models{i} = model;
        oneminute_target(i) = oneminute;
        max_target(i) = max;
        i=i+1;
    end
    k=k+1;
end
close(h1);
    function random_p = get_random(variable, num)
        dist = fitdist(variable,'gamma');
        random_p = random(dist, num,1);
    end
end
```

```
function [oneminute_target,max_target] = modelTargets(model)

    f = @(thebolus)maxObjectiveFunction(thebolus, model);
    max_target = fminbnd(f,.2,6);
    f = @(thebolus)twominuteObjectiveFunction(thebolus, model, 1);
    oneminute_target = fminbnd(f,.2,max_target);

    function error = maxObjectiveFunction(thebolus, model)
        target = model.target_conc;
        t=0:1/60:6;
        u=zeros(size(t));
        u(1) = 60*thebolus;
        conc = lsim(model.sys,u,t);
        error = abs(conc(end)-target);
    end

    function error = twominuteObjectiveFunction(thebolus, model, duration)
        target = model.target_conc;
        t=0:1/60:6;
        u=zeros(size(t));
        u(1) = 60*thebolus;
        conc = lsim(model.sys,u,t);
        a = conc>=target;
        if sum(a) < 3
            error = 2 - sum(a)/60;
        else
            b = find(a==1,1,'first') +[-1 0];
            c = find(a==1,1,'last')+ [0 1];
            uptime = interp1(conc(b),t(b),target);
            if (any(c>length(a)))
                downtime = t(end) + 1/60;
            else
                downtime = interp1(conc(c),t(c),target);
            end
            rawerror = downtime - uptime - duration;
            error = abs(rawerror);
        end
    end

end
```
